# The Heterogeneous Solution Ensemble of the DEAD-box Protein Dhh1 Reveals a Modular Architecture

**DOI:** 10.64898/2025.12.08.692948

**Authors:** Davide Erba, Edoardo Fatti, Matteo Paloni, Karsten Weis, Pau Bernadó, Alessandro Barducci

## Abstract

DEAD-box proteins such as Dhh1 play essential roles in RNA metabolism and the formation of biomolecular condensates, with a modular architecture comprising folded domains and disordered regions. To elucidate how this architecture shapes conformational dynamics in solution, we combined solution scattering experiments and multi-scale simulations on core and full-length constructs. Enhanced-sampling simulations captured a dynamic ensemble of core conformations stabilized by transient interdomain contacts that underpin functional regulation. Coarse-grained modelling revealed that disordered tails behave as independent modules exerting minimal influence on core dynamics. This integrated approach reveals a modular organization balancing structural heterogeneity and functional specificity, providing a framework for studying DEAD-box proteins in phase separation.

## Introduction

Helicases are essential enzymes that couple ATP binding and hydrolysis to the remodelling and unwinding of nucleic acids, participating in numerous aspects of RNA and DNA metabolism Fairman-Williams et al. (2010); Jankowsky (2011). Among them, DEAD-box proteins (DBPs), which form the largest subgroup within the SF2 superfamily, are defined by a conserved D-E-A-D motif critical for ATP hydrolysis and a core architecture comprising two RecA-like domains connected by a linker Weis (2021); Banroques et al. (2011). These enzymes mediate local and sequence-independent changes in RNA structure, underlining their diverse roles from transcription to RNA degradation and storage Weis and Hondele (2022); Linder and Jankowsky (2011).

A hallmark of DBPs is their structure: a folded core divided into two RecA-like domains flanked by flexible tails (figure 1A). This architecture enables a conformational cycle involving ATP and RNA binding, hydrolysis, and RNA release, yet many mechanistic aspects remain poorly resolved. Recent works have highlighted DEAD-box proteins as key scaffolds of membraneless organelles *in vivo*, where they participate in the formation and regulation of biomolecular condensates Hondele et al. (2019); Mugler et al. (2016). *In vitro* studies have revealed that the intrinsic structural features of these proteins, particularly their disordered tails, play a crucial role in promoting phase separation Hondele et al. (2019). Additionally, environmental conditions such as pH, salt and RNA concentration strongly modulate their phase separation behaviour, emphasizing the interplay of molecular architecture and cellular context in driving condensate assembly Hondele et al. (2019).

**Figure 1.**
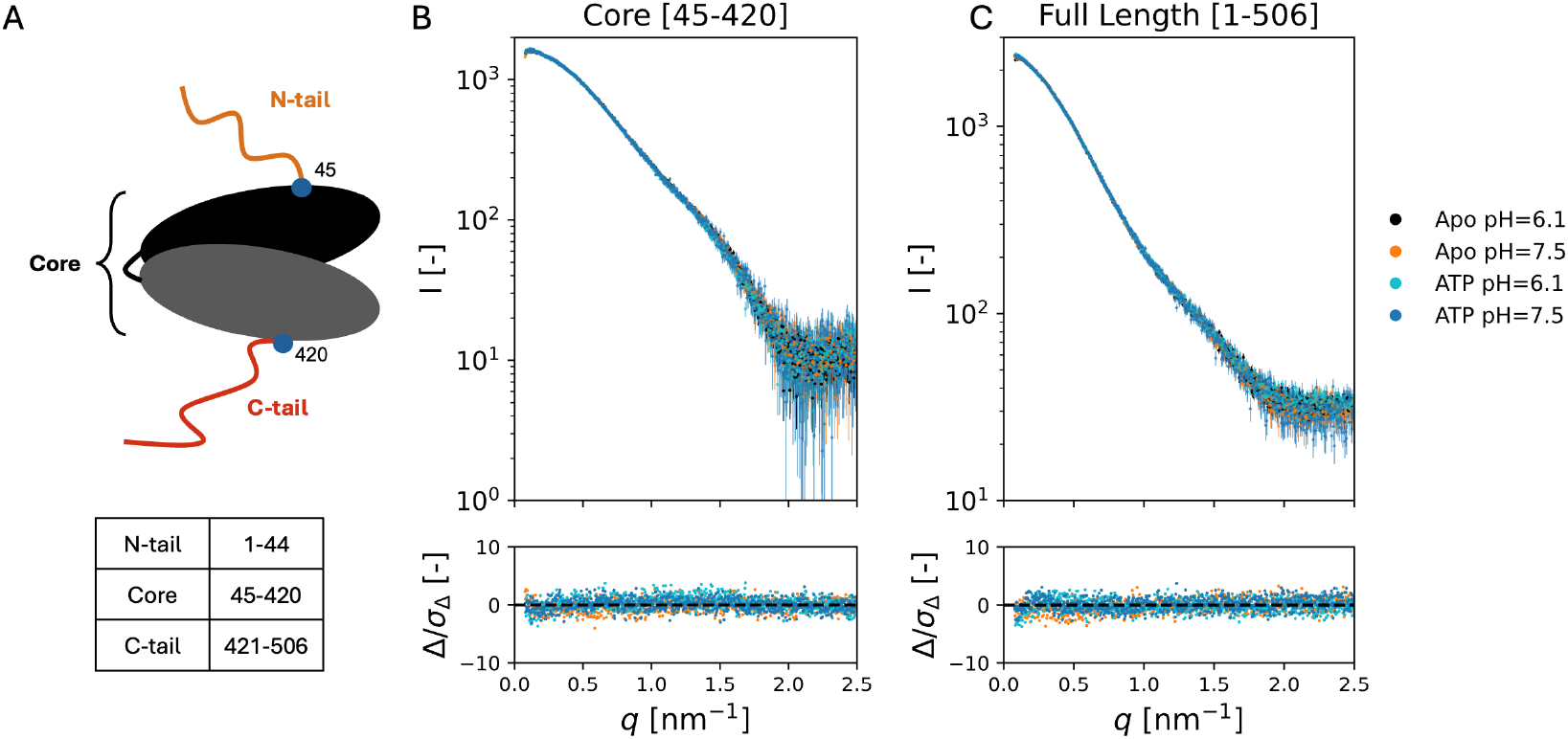
**A.** *Top*: Cartoon of prototypical DEAD-box protein, characterised by a folded core divided into two RecA domains connected by a linker and flanked by two disordered tails. *Bottom*: Definition of domains for Dhh1. **B**. SAXS measurements (top) performed on the core sequence of Dhh1, corresponding to the sequence 45-420, together with the distribution of residuals (bottom), show that pH variation and ATP addition do not affect the observed profiles. **C**. The FL of Dhh1 is insensitive to pH changes and ATP addition, too. **B**,**C**. Given the similarity of the profiles across conditions, the ones obtained in Apo conditions at pH=6.1 will be showed as representative of the experiments.

Dhh1 is a DEAD-box protein from *Saccharomyces cerevisiae* that acts in translational repression and mRNA storage Cheng et al. (2005); Linder and Jankowsky (2011). Unlike many DEAD-box proteins, Dhh1 can bind RNA independently of ATP, and its ATPase activity is modulated by cofactors Cheng et al. (2005); Sharif et al. (2013). The structure of Dhh1 is prototypical of DEAD-box proteins, with a folded core spanning residues 45-420 flanked by disordered N-tail (corresponding to residues 1-44) and the C-tail (421-506), as shown in figure 1A. Currently, only one crystal structure, obtained in Apo conditions (PDB code 1S2M Cheng et al. (2005), in gray in figure 2A), is available. This structure reveals unique interdomain contacts in the absence of any ligand that were further explored by mutagenesis. Concretely, R89A, K91A and R345A mutants showed altered RNA affinity and ATPase activity Cheng et al. (2005); Dutta et al. (2011). Predicted models diverge, with AlphaFold3 Abramson et al. (2024) proposing a compact conformation distinct from the available experimental structure (in red in figure 2A), but resembling conformations observed for other DBPs bound to RNA and nucleotides Andersen et al. (2006); Ngo et al. (2019).

**Figure 2.**
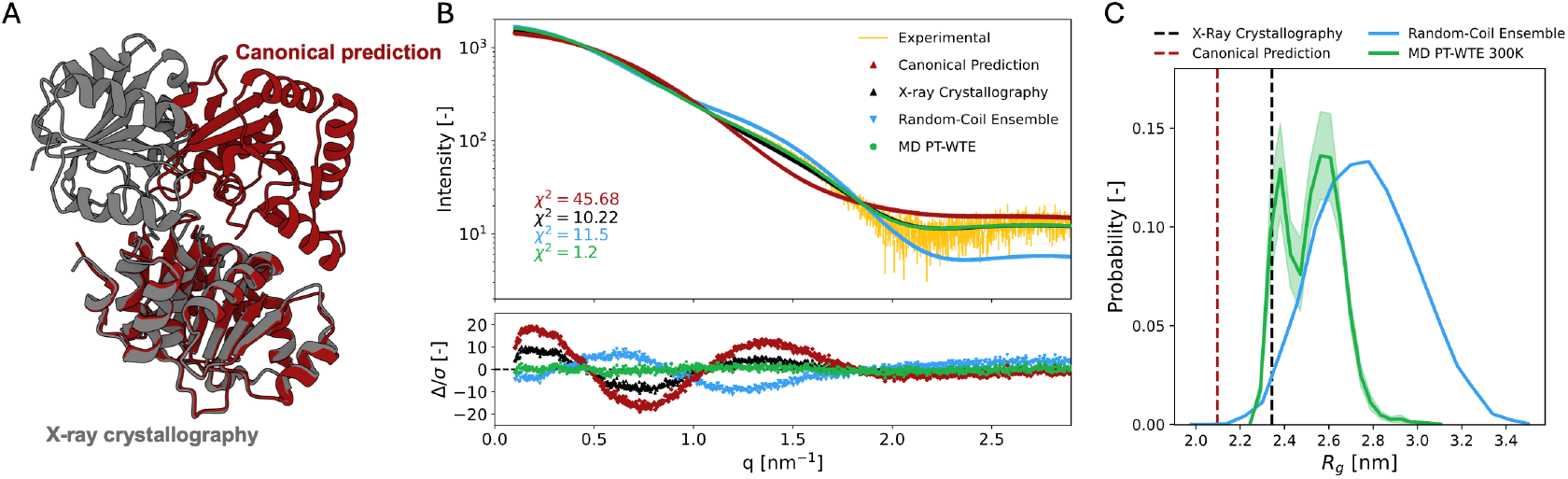
**A**: Snapshots of the crystallographic X-ray structure 1S2M (gray) and of the canonical rearrangement (red), after alignment on the RecA1 to highlight the rearrangement of the RecA2. **B**: Comparison of the SAXS experimental profile (yellow) with those corresponding to the crystallographic structure 1S2M (black), the canonical rearrangement (red), that obtained from an ensemble of 10^4^ structures with random-coil linker (light blue), and that from the production trajectory at 300K (green). **C**: Distribution of the radius of gyration (*R*_*g*_ ) obtained from the MD production trajectory. The curve represents the average over five independent 200 ns segments of the trajectory, which yielded consistent distributions. The bimodal shape indicates that the Dhh1 core samples both compact conformers (vertical dashed lines), compatible with the crystallographic structure, and more extended conformations that remain significantly smaller than those accessible in the random-coil model (blue)

Despite these advances, the solution behaviour and functional conformational states of Dhh1 remain largely unexplored. The available structure provides only a static view under potentially non-physiological conditions, leaving unresolved the extent of its conformational plasticity and the role of the tails in Dhh1 dynamics. In order to address these points, we applied a combined approach integrating Size Exclusion Chromatography online with Small-Angle X-ray Scattering (SEC-SAXS) and molecular dynamics (MD) simulations. On the one hand, for the core construct of Dhh1, we coupled SAXS to enhanced atomistic simulations, taking advantage of the Parallel Tempering in the Well-Tempered Ensemble (PT-WTE) Prakash et al. (2011); Barducci et al. (2015). On the other hand, for the full-length (FL) construct, we combined SAXS to coarse-grained, one-bead-per-residue simulations. The overall model emerging from our synergistic approach describes Dhh1 as a modular protein, where the tails and the core act as independent and decoupled compartments, providing the structural bases of its function.

## Results

### SAXS Defines the Global Solution Behaviour of Dhh1 Across Conditions

To dissect the structural contributions of the folded RecA domains and the disordered tails, we studied two constructs of Dhh1: a core construct comprising residues 45–420 and the full-length (FL) protein. We characterised the solution conformation of both constructs by SEC-SAXS, which avoids the contribution from aggregates and ensures that data from the monomeric species are measured. For each construct, we measured profiles at two pH values, 6.1 and 7.5, which are known to yield different condensation propensities Hondele et al. (2019); Mugler et al. (2016), and in the absence or presence of ATP, resulting in four measurements per construct.

For the core construct, the experimental profiles seemed insensitive to the changes in pH and the presence of ATP. The structural parameters obtained from Guinier and *p*(*r*) analyses (Table 1) yielded *R*_*g*_ values in the range 2.62-2.64 nm and a maximum particle distance *D*_*max*_ of 8.8-9.0 nm. After proper rescaling of each profile against the Apo pH 6.1 dataset, the residuals were centred around zero and *ω*^2^ values remained low. In the normalised Kratky representation (figure S1) all curves showed the expected bell-shaped profile characteristic of compact molecules.

A similar behaviour was observed for the FL construct. Guinier and *p*(*r*) analyses (Table 2) yielded *R*_*g*_ values between 3.38 and 3.41 nm, with *D*_*max*_ around 13.5 nm. The four profiles in figure 1C overlapped well, with low and unbiased residuals when compared to the Apo pH 6.1 profile. The normalised Kratky plot (figure S2) displayed a pronounced peak at low *qR*_*g*_, as expected for a folded core, followed by a rising tail at higher *qR*_*g*_ values, consistent with the presence of disordered termini. Overall, these results indicate that the global features of Dhh1, for both the core and FL constructs, are conserved across the tested conditions. Neither pH variation nor ATP addition produces detectable large-scale structural rearrangements in solution under our experimental conditions. In particular, the absence of structural changes upon ATP addition is in line with previous biochemical and structural work on DEAD-box proteins; indeed, for DEAD-box ATPases, ATP binding alone does not stabilise the closed RecA-domain conformation and typically requires cooperative RNA binding to populate this compact state Cordin et al. (2006); Linder and Jankowsky (2011); Liu et al. (2008). Like-wise, the lack of pH dependence in our SAXS profiles shows that the monomeric architecture of Dhh1 remains unchanged between pH 6.1 and 7.5. This is counter-intuitive, given the known pH sensitivity of Dhh1 condensation, which arises in the presence of RNA Hondele et al. (2019); Mugler et al. (2016). Together, these observations suggest that pH may primarily modulate intermolecular interactions rather than altering the global conformation of isolated Dhh1, and that Dhh1 requires RNA to proficiently bind ATP.

### Enhanced-Sampling Atomistic MD Reveals a Heterogeneous Core Ensemble Consistent with SAXS Data

To evaluate whether available structural models could explain the SAXS profile of the Dhh1 core, we compared the experimental curve with two compact conformations shown in figure 2A. The first is the crystallographic Apo structure 1S2M Cheng et al. (2005). The second represents the canonical closed arrangement adopted by ligand-bound DEAD-box proteins, where the two RecA domains form the ATP- and RNA-binding pockets, as observed in several structures Andersen et al. (2006); Ngo et al. (2019). When compared to the experimental SAXS data collected under Apo conditions at pH 6.1, neither model reproduces the observed profile, resulting in high *χ*^2^ values and systematic deviations in the point-by-point residual profile (figure 2B). This indicates that no available compact structure can account for the behaviour of Dhh1 in solution.

Because structural models were insufficient to explain the SAXS profile, we turned to molecular dynamics (MD) simulations and used PT-WTE, an enhanced-sampling method that combines replica exchange with a metadynamics bias on the potential energy and allows efficient exploration of alternative arrangements of the two RecA domains. The *in silico* SAXS profile computed from the 300 K production trajectory shows excellent agreement with the experimental curve, with *χ*^2^ = 1.2 and residuals distributed symmetrically around zero (figure 2B). This agreement suggests that the ensemble sampled in the simulation provides an accurate description of the solution-state structure of the Dhh1 core.

Then we examined the distribution of the radius of gyration (*R*_*g*_) across the trajectory to characterise the underlying heterogeneity (figure 2C). The resulting distribution was bimodal. The first mode contains compact conformers whose size is compatible with the crystallographic structure, yet conformations with an *R*_*g*_ comparable to the canonical closed DEAD-box arrangement are not sampled, where the second mode corresponds to more extended conformations.

To test whether simple linker flexibility could reproduce the SAXS profile, we generated a control ensemble using RanCh Franke et al. (2025), in which the two RecA domains are held fixed in their crystallographic structures but connected by a linker that can freely rearrange. The average scattering profile of this ensemble does not satisfactorily reproduce the experimental data and shows a systematic loss of intensity at small *q* values (figure 2B, light blue), indicating that these conformations are too extended. This result shows that conformational heterogeneity alone is not sufficient to explain the data. The ensemble must include specific, structured domain arrangements, such as those sampled in the MD simulations, in order to match the SAXS profile. Taken together, these results show that the Dhh1 core samples a heterogeneous ensemble of compact and moderately extended conformations in solution, and that this ensemble is required to reproduce the SAXS data. Static models, whether crystallographic, canonical, or random, cannot account for the experimental profile on their own.

### A Dynamic Conformational Landscape with Frequent Interdomain Contacts Reconciles Ensemble Heterogeneity with Functional Mutagenesis Data

Having established that the MD ensemble reproduces the experimental SAXS profile, we next characterised its conformational organisation. The free-energy surface (FES, figure 3A) was estimated in the two-dimensional space defined by *R*_*g*_ and the RMSD computed against the experimental structure 1S2M. The FES displays several local minima of comparable depth, one of which corresponds to conformers similar to the crystallographic structure, as indicated by the low RMSD. To better characterise the conformations associated with these minima, we performed clustering on the production trajectory by retaining 10^5^ equally spaced frames and identified a representative structure (centroid) for each cluster (figure 3B and Supplementary figure S3). The clustering procedure revealed several distinct clusters. In particular, the most populated one corresponds to a set of frames compatible with the crystallographic structure and consists of about 12% of the trajectory frames. This shows that the 1S2M conformation is present in solution but does not dominate the ensemble. In this sense, the X-ray model provides a structurally precise but partial snapshot of a broader and dynamic conformational landscape. Other clusters populate more extended or differently oriented states, in agreement with the bimodal *R*_*g*_ distribution described above. Visual inspection of the centroids of the most populated clusters (figure 3B and supplementary figure S4) supports a Dhh1 model in which linker flexibility allows different rearrangements of the two RecA domains.

**Figure 3.**
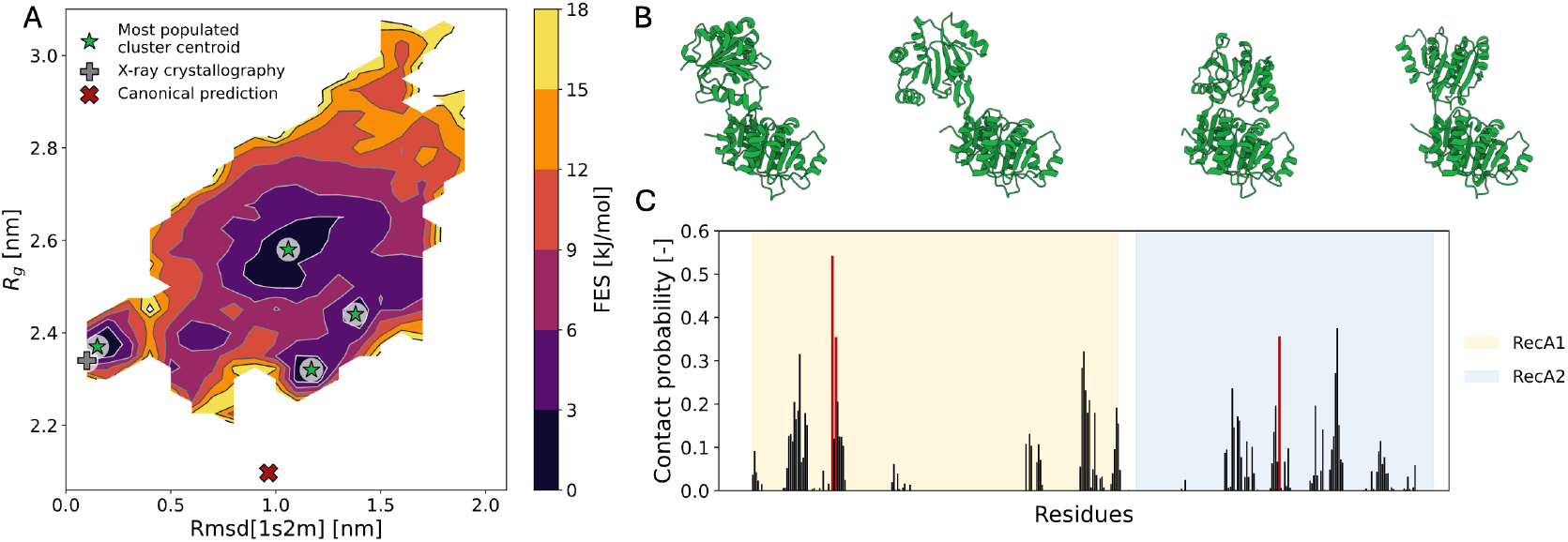
**A**: FES computed in the *R*_*g*_ /RMSD space highlights the diversity of the Dhh1 core conformers. In particular, the crystallographic structure (gray cross) is found in one of the minima, revealing its importance, whereas the canonical conformation (red cross) lies in a region of the FES not explored by the trajectory. The stars indicate the position of the static structures in the 2D space. **B**: The centroids of the four most-populated clusters, whose position in the FES is indicated by green stars, display different arrangements of the two RecA domains. **C**: Per-residue interdomain contact probability highlights that three of the four most interacting residues are ARG89, LYS91 and ARG345. These residues have been experimentally identified to form interdomain contacts that affect Dhh1 activity.

To characterise the differences between the four most populated clusters, we computed the average interdomain C_*α*_ contact maps (supplementary figure S5). The overlapping maps reveal that, on average, each conformational cluster is characterised by distinct interdomain interacting regions. In particular, some regions of RecA1 exhibit interaction propensities with multiple regions of RecA2, consistent with linker flexibility that permits a wide range of interdomain arrangements, and thus contacts.

The reduced activity of Dhh1 has been strictly associated with the compact conformation observed in the crystal structure 1S2M Cheng et al. (2005). The physiological relevance of this conformer was further supported by mutagenesis experiments that targeted residues involved in interdomain contacts, in particular ARG89, LYS91 and ARG345 Cheng et al. (2005); Dutta et al. (2011). However, our SAXS-consistent model contains only about 12% of frames similar to the crystallographic structure, which at first sight seems at odds with the strong functional effects of these mutations. To investigate this apparent contradiction, we estimated the probability of each residue to interact with the opposite RecA domain (figure 3C). This analysis reveals that both RecA domains contain regions with a nonzero probability of forming interdomain contacts. Strikingly, three of the four residues with the highest interdomain contact probabilities, R89, K91 and R345 (red bars in figure 3C), are precisely those whose mutation enhances Dhh1 activity Cheng et al. (2005); Dutta et al. (2011). Our results therefore suggest that these mutations alter the conformational ensemble by perturbing a broad and dynamic network of interdomain contacts, rather than only the specific interface stabilised in the crystallographic structure.

Overall, these results support a model in which the Dhh1 linker rearranges the two RecA domains, allowing conformers with different sizes and orientations than the crystallographic structure, which still remains the most populated conformational state of Dhh1. At the same time, a set of recurrent interdomain contacts is preserved across the ensemble, providing a structural basis that reconciles the heterogeneous conformational landscape with the functional relevance of residues identified in mutagenesis experiments.

### Coarse-Grained Simulations Reveal a Modular Architecture with Disordered Tails Decoupled from Core Dynamics

To extend our structural analysis beyond the isolated core and describe the behaviour of full-length Dhh1 (FL) in solution, we turned to coarse-grained (CG) modelling. Full-length Dhh1 is too large and flexible to be sampled exhaustively by explicit-solvent atomistic MD, so we used CALVADOS Tesei et al. (2021); Tesei and Lindorff-Larsen (2023); Cao et al. (2024), a 1-bead-per-residue representation that accurately reproduces the conformational ensembles of intrinsically disordered proteins. Since this model does not preserve secondary structure, each RecA domain was kept in its experimental fold by intra-domain restraints, and we evaluated how different assumptions about interdomain contacts and overall core flexibility affected agreement with the FL SAXS profile.

As a first test, we generated an ensemble in which flexible tails were added to the crystallographic core. The resulting SAXS curve differs markedly from the experimental profile, with large residuals indicating that this ensemble remains too compact (figure 4B, cyan). Thus, simply adding disordered tails to a rigid core is not sufficient to reproduce the behaviour of full-length Dhh1 in solution.

**Figure 4.**
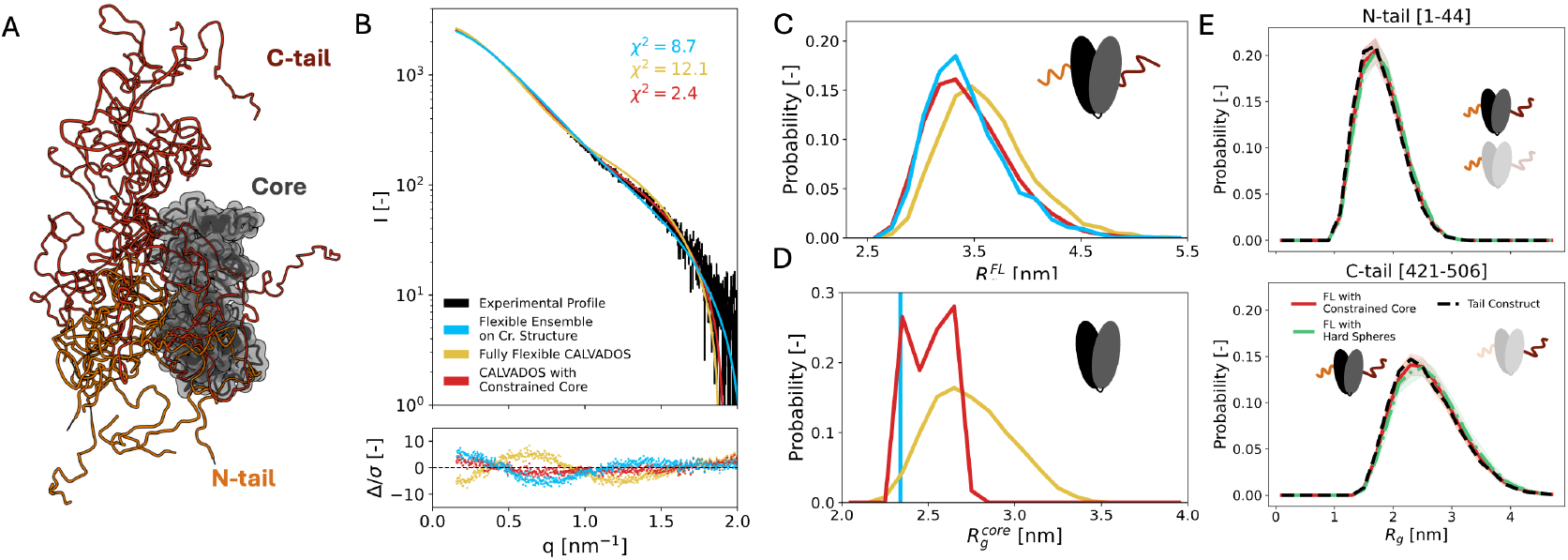
**A.** Snapshots of Dhh1 FL from CALVADOS simulations **B**. Experimental SAXS profile in Apo conditions at pH 6.1 is compared to the *in silico* profiles computed from CALVADOS simulations. Blue points indicate the disordered ensemble built adding tails to the core in 1S2M conformation, yellow points indicate CALVADOS simulations where the interdomain contacts are not restrained, and therefore the core is flexible, and red points indicate simulations where the core is restrained to respect the results obtained with atomistic simulations (labelled as Clustered Core). **C**. The distribution of the FL construct is bell-shaped in all three cases, but the presence of restraints shifts the mean value from 3.5 nm to 3.3 nm. Notably, the relative flexibility allowed by restraints does not affect the character of the FL distribution, but drastically improves the quality of the SAXS comparison **D**: Comparison of the core *R*_*g*_ distribution show that the Full-Flex model has a bell-shaped distribution compatible with an ensemble of structures with random-coil linker. Conversely, the restrained simulations adapt their core size distribution to the one observed at atomistic resolution. **E**. Agreement between *R*_*g*_ distributions of the N-tail (left) and C-tail (right) computed with restrained simulations and with a hard-sphere core suggest that the presence of the core does not influence the behaviour of both tails.

We next examined whether the presence of the tails could unlock the core and generate a more extended ensemble. To this end, we performed fully flexible CALVADOS simulations in which the RecA domains were allowed to rearrange freely. This model yields a SAXS profile that underestimates the intensity at low *q*-values, consistent with an ensemble that is too expanded (figure 4B, yellow). The corresponding core *R*_*g*_ distribution includes conformations extending beyond 3.5 nm (figure 4D), in clear disagreement with both SAXS and atomistic MD. These results show that unrestrained CALVADOS simulations overestimate core flexibility and cannot account for the experimental data.

Finally, we introduced restraints that restricted the RecA-RecA arrangement to the conformational ensemble previously obtained for the isolated core construct. This model yields a SAXS profile in excellent agreement with the experimental full-length curve, with small and structureless point-by-point residuals (figure 4B, red). The full-length *R*_*g*_ distribution remains broad in all cases (figure 4C), but the restrained-core ensemble is slightly more compact than the fully flexible model. Importantly, its core *R*_*g*_ distribution recovers the bimodal behaviour observed in the atomistic ensemble (figure 4D), confirming that the CG model faithfully propagates key features of the MD-derived conformational landscape. The fact that this ensemble reproduces the full-length SAXS data already suggests that the disordered tails do not appreciably perturb core dynamics.

We then characterised the behaviour of the Nand C-terminal tails within this FL ensemble. Intra-tail contact maps do not reveal persistent interaction patterns (supplementary figure S6), consistent with their intrinsically disordered nature. The *R*_*g*_ distributions of both tails are unimodal (figure 4E), with mean values that scale with sequence length. These distributions are nearly identical when the core is represented as a hard sphere or when the tails are simulated alone, indicating that tail behaviour is largely insensitive to the state of the core. Additional analyses of tail-core contacts support this picture: interactions are sparse, highly transient, localised near the attachment points, and do not depend on the specific RecA-RecA arrangement (supplementary figure S7).

Taken together, these results highlight the modular organisation of Dhh1. The structured core is best described by the heterogeneous ensemble observed in isolation, and its conformational landscape is preserved in the full-length protein, whereas the disordered tails behave as independent units whose structural properties do not depend on the core. This modular architecture provides a coherent framework for understanding the solution behaviour of full-length Dhh1.

## Conclusion

This study provides an integrated molecular description of the DEAD-box protein Dhh1 by combining SEC-SAXS measurements with atomistic and coarse-grained simulations. Our results reveal that, rather than adopting a single dominant conformation, the Dhh1 core samples a heterogeneous ensemble in solution. This ensemble is shaped by linker flexibility and consists of multiple interdomain arrangements, of which the crystallographic Apo structure represents only one accessible state.

Despite this structural heterogeneity, the ensemble displays a set of recurrent interdomain contacts. Several of the residues that participate most frequently in these contacts, such as ARR89, LYS91 and ARG345, have long been known to modulate ATPase and RNA-binding activity when mutated Cheng et al. (2005); Dutta et al. (2011). Our results therefore reconcile the heterogeneous solution ensemble with classical mutagenesis data: these residues are functionally important not because they stabilise a single crystallographic interface, but rather because they contribute to a broader and dynamic interaction network that recurs across many conformational states.

Extending our analysis to the FL protein, CG simulations demonstrate that the behaviour of Dhh1 is fundamentally modular. The disordered N- and C-terminal tails behave as flexible, independent segments whose conformational properties are largely insensitive to the specific arrangement of the core domains. Accurate reproduction of the FL SAXS profile requires preserving the core ensemble identified in atomistic simulations, indicating that the tails do not perturb core dynamics. This modular architecture offers a natural explanation for the observation that pH-dependent condensation requires RNA and cofactors: monomeric Dhh1 remains structurally stable in solution, while the disordered tails likely contribute to intermolecular interactions that drive condensate formation.

Together, these findings highlight the synergy between SAXS and multi-scale simulations for dissecting the conformational landscapes of multi-domain RNA-binding proteins. Dhh1 emerges as a highly modular molecule in which a dynamically regulated core is coupled to intrinsically disordered tails that tune interaction propensity without altering core structure. This framework provides a mechanistic basis for understanding DEAD-box ATPases in both their enzymatic roles and their contributions to RNA–protein condensates, and offers general principles applicable to the broader family of helicase proteins.

## Methods

### Expression and Purification of Dhh1

Dhh1FL (pKW5049), and Dhh145-420(pKW5113) were expressed in *Escherichia coli* Rosetta (DE3) (Novagen) as 6xHis-TEV tagged proteins using an auto-induction medium. Cells were grown at 37 °C in 1 L ZY complete medium to an OD600 of 0.7, then the temperature was lowered to 20 °C and growth continued for 20 h at 220 rpm. Cells were harvested, washed in cold PBS and the cell pellet was flash frozen in liquid nitrogen and stored at -20 °C until use. The pellet was dissolved in 10 mL/g of Lysis Buffer (20 mM HEPES-KOH pH 7.7, 500 mM KCl, 5 mM MgCl_2_, 0.2% IGEPAL CA-630, 10 mM Imidazole, 2 mM β-mercaptoethanol) supplemented with 0.5 mg/mL Lysozyme (PanReac AppliChem) and 0.01 mg/mL DNAse I (PanReac AppliChem), and mechanically disrupted by high pressure homogenizer (Emulsiflex C5, Avestin). The lysate was centrifuged at 20,000xg (SS-34 fixed angle rotor, Sorvall) and the supernatant was filtered through a 0.45 *μ*m PES membrane filter (Sarstedt) and incubated with NiNTA beads (Qiagen). Beads were washed with 5 column volumes of Detergent Buffer (20 mM HEPES-KOH pH 7.7, 500 mM KCl, 5 mM MgCl_2_, 0.2% IGEPAL CA-630, 10 mM Imidazole, 2 mM β-mercaptoethanol, 5% Glycerol), High Salt Buffer (20 mM HEPES-KOH pH 7.7, 1.5 M KCl, 5 mM MgCl_2_, 10 mM Imidazole, 2 mM β-mercaptoethanol, 5% Glycerol), Imidazole Buffer (20 mM HEPES-KOH pH 7.7, 1.5 M KCl, 5 mM MgCl_2_, 20 mM Imidazole, 2 mM β-mercaptoethanol, 5% Glycerol), and eluted in 2.5 column volumes of Elution Buffer (20 mM HEPES-KOH pH 7.7, 500 mM KCl, 5 mM MgCl_2_, 330 mM Imidazole, 2 mM β-mercaptoethanol, 5% Glycerol). The elution buffer was exchanged to Imidazole Buffer by passing the proteins through a PD-10 column (GE Healthcare) and incubated overnight at 10 °C with 6xHis-TEV protease. The eluate was passed again through Ni-NTA beads to remove uncleaved proteins and the 6xHis-TEV protease, concentrated with centrifugal filter units (Millipore) and further purified by size exclusion chromatography on a Superdex 200 16/600 column (GE Healthcare) in SEC Buffer (20 mM HEPES-KOH pH 7.7, 500 mM KCl, 5 mM MgCl_2_, 1 mM DTT, 10% Glycerol) using an AKTA pure system (GE Healthcare). Positive fractions were pooled, further concentrated and final purity was assessed by SDS-PAGE and Coomassie stain (Instant Blue®, Abcam). Aliquots were flash frozen and stored at -80 °C.

### SAXS Measurements

In all SAXS measurements, the protein was at a concentration of approximately 75 *μ*M, which corresponds to approximately 3 mg/mL sample for the core sequence and 4.5 mg/mL for the FL. Passing the sample through a 24 mL S200 SEC column dilutes the protein roughly ten-fold. The buffer of SAXS measurements performed in absence of nucleotide contained 20 mM HEPESKOH pH 7.7, 300 mM KCl, 5 mM MgCl_2_, 5% glycerol and 1 mM DTT. The buffer of measurements in presence of ADP/ATP contained 20 mM PIPES-KOH pH 6.1 or 7.5, 300 mM KCl, 5 mM MgCl_2_, 2,5% glycerol and 1 mM DTT. After equilibration, the measurements were performed for one hour, collecting 3000 individual frames, corresponding to a frame every 1.2s.

Frame selection and background subtraction from SEC-SAXS measurements were performed using Chromixs (ATSAS 3.2.1) Manalastas-Cantos et al. (2021), while raw data analysis (Guinier and *p*(*r*) analysis) were performed with Primus (ATSAS 4.0.0) Franke et al. (2025).

#### In silico SAXS Profile Computation and Handling

*In silico* scattering profiles for atomistic conformations, both static structures and the MD trajectory were computed using Crysol from ATSAS 4.0.0 Franke et al. (2025) with the default options, 20 spherical harmonics and Fibonacci grid of order 17, and eventually adding the explicit-hydrogens option to solve possible clashes due to atom names. Scattering profiles for one-bead-per-residue were computed using the PLUMED extension hySAS Ballabio et al. (2023) using *C*_*α*_ centred beads.

#### Agreement Estimation

Reduced *χ*^2^ was computed to assess the agreement between profiles defined on the same range of *N* wave vectors *q*. If both profiles (*I*_1_, *I*_2_) are characterised by experimental error, the definition is

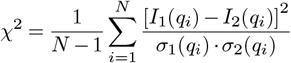

while, if only *I*_1_ has experimental error, it becomes:

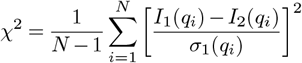

Rescaling and subtraction of a profile *I*_2_ against *I*_1_ was performed using an in-house Python script that searches for the *γ* parameters that minimises the *χ*^2^ 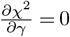.

Residuals between a profile characterised by error (*I*_1_, *σ*_1_) and a profile without errors (*I*_2_, rescaled by *γ*) were defined as

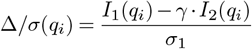

Conversely, residuals between two profiles characterised by errors (*I*_1_, *σ*_1_ and *I*_2_, *σ*_2_, *γ*) were defined as

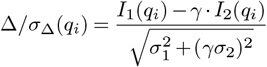

### Atomistic Simulations

Atomistic PT-WTE simulations were performed using GROMACS 2021.4 Abraham et al. (2015) patched with Plumed 2.8.0 Tribello et al. (2014); Bonomi et al. (2019). We used 32 replicas, and determined temperatures following equations (4) and (5) in Prakash et al. (2011), selecting *T*_0_ = 300K and *T*_1_ = 305K, so that the highest temperature is *T*_31_ = 542.5K.

The initial configuration of Dhh1 was the experimental structure 1S2M, modelled with the DES-Amber force field with charge rescale Tucker et al. (2022). The structure was solvated using TIP4P-D Henriques and Skepö (2016) in a rhombic dodecahedron box volume of approximately 1272nm^3^, and then neutralized with NaCl, adjusting salt concentration to 0.3M. After energy minimisation, a first 100 ps equilibration was run in the NVT ensemble at *T* = *T*_0_, followed by a 100 ps NPT simulation at *T* = *T*_0_ and at *P* = 1bar, controlling temperature and pressure respectively by the v-rescale Bussi et al. (2007) and the Parrinello-Rahman Parrinello and Rahman (1981) schemes. Then, for each temperature, we performed 1 ns simulation in the NVT ensemble, followed by a 5 ns WTE simulation Bonomi and Parrinello (2010) deposing Gaussian bias on the energy of the system every ps, with Gaussian width of 500kJ/mol, and height of 2kJ/mol with a biasfactor *b* = 10. Finally, we performed 1.3 *μs* PT-WTE simulation, with odd replica exchange attempted every 10 ns Barducci et al. (2015) and bias deposited on the potential energy every ps.

For all simulations, we applied periodic boundary conditions, and long-range electrostatic interactions were evaluated by using the particle mesh Ewald algorithm Darden et al. (1993); Essmann et al. (1995), with a 1nm cut-off for the real space interactions, and van der Waals interactions were computed by using the same cut-off distance. The lengths of bonds involving hydrogen atoms were constrained using LINCS algorithm Hess et al. (1997); Hess (2008), allowing for a time step of 2fs to integrate the equations of motion. To prevent unfolding of Dhh1 structured domains, RMSD restraints were applied to each RecA domain based on their coordinates in the experimental structure, by using PLUMED Tribello et al. (2014); Bonomi et al. (2019) `UPPER_WALLS` on single domain’s RMSD with a harmonic constant of 50000kJ/mol.

Atomistic simulations were analysed using a mix of GROMACS Abraham et al. (2015), PLUMED Tribello et al. (2014); Bonomi et al. (2019) and MDAnalysis Michaud-Agrawal et al. (2011) functions and modules.

Conformational clustering was performed using the GROMACS `cluster` command with the algorithm proposed by Daura Daura et al. (1999) on a trajectory consisting of 10^4^ equally spaced frames and with a neighbouring distance cut-off of 0.35 nm. Radius of gyration *R*_*g*_ was computed on C_*α*_ atoms using PLUMED `GYRATION` function. Root mean square deviation RMSD was computed on C_*α*_ atoms after alignment of RecA1 on the reference structure with PLUMED `RMSD` function. Interdomain contacts were computed using the MDAnalysis `distance` function over C_*α*_ atoms (with a cutoff distance *r*_*c*_ = 0.8 nm) or heavy atoms (*r*_*c*_ = 0.5 nm).

### Coarse-Grained Simulations

One-bead-per-residue simulations of FL Dhh1 were performed using the CALVADOS3 force field Tesei et al. (2021); Tesei and Lindorff-Larsen (2023); Cao et al. (2024), taking advantage of the python simulation framework. In all simulations, residues in folded (or restrained) regions were represented by a bead positioned on their centre of mass, while residues in flexible regions by a bead centred on the C_*α*_ atom.

Flexible CALVADOS simulations were performed by constraining each single core with an elastic network using a cut-off distance of 0.9 nm. Three independent replicas of 500 ns were performed, yielding 1.5 · 10^4^ total frames to analyse.

Simulations with restrained core were performed by adding a supplementary elastic network to fix the conformation of the core (corresponding to residues 45-420). Each starting conformer was built by adding disordered tails to the centroids of the most populated clusters, as obtained from previous conformational clustering on atomistic simulations. For each centroid, three independent replicas of 500 ns were performed, saving one frame each 500 ps.

CALVADOS simulations were analysed using MDAnalysis Michaud-Agrawal et al. (2011) functions and modules. Radius of gyration *R*_*g*_ was computed using the MDAnalysis `radius_of_gyration` function, while interdomain contacts were computed using the `distance` function.

### Generation of Random Coil Structures

Random coil ensembles of conformations were generated using RanCh (ATSAS 4.0.0) Franke et al. (2025). Each conformer was generated by constraining each RecA in the conformation of the crystallographic structure, fixing RecA1 in space and allowing RecA2 to rearrange in space after generating a random coil linker.

## Notes

### Competing Interest Statement

The authors have declared no competing interest.

### Funding

- Swiss National Science Foundation: CRSII5_193740

