## Supplementary tables and figures for "The Heterogeneous Solution Ensemble of the DEAD-box Protein Dhh1 Reveals a Modular Architecture"

### Supplementary Information

**Table 1.** Raw data analysis from the SAXS profiles obtained for the core sequence of Dhh1 by varying pH and inserting ATP.

| pH | Buffer | $R_g$ [nm] | $D_{max}$ [nm] | $\chi^2$ [Apo 6.1] |
|---|---|---|---|---|
| 6.1 | Apo | 2.62±0.01 | 8.8 | - |
| 6.1 | ATP | 2.64±0.01 | 9.0 | 1.54 |
| 7.5 | Apo | 2.64±0.01 | 8.8 | 1.22 |
| 7.5 | ATP | 2.62±0.01 | 9.0 | 1.31 |

**Table 2.** Raw data analysis from the SAXS profiles obtained for the FL sequence of Dhh1 by varying pH and inserting ATP.

| pH | Buffer | $R_g$ [nm] | $D_{max}$ [nm] | $\chi^2$ [Apo 6.1] |
|---|---|---|---|---|
| 6.1 | Apo | 3.38±0.01 | 13.5 | - |
| 6.1 | ATP | 3.41±0.01 | 13.0 | 1.27 |
| 7.5 | Apo | 3.40±0.01 | 13.7 | 1.50 |
| 7.5 | ATP | 3.41±0.01 | 13.3 | 1.49 |

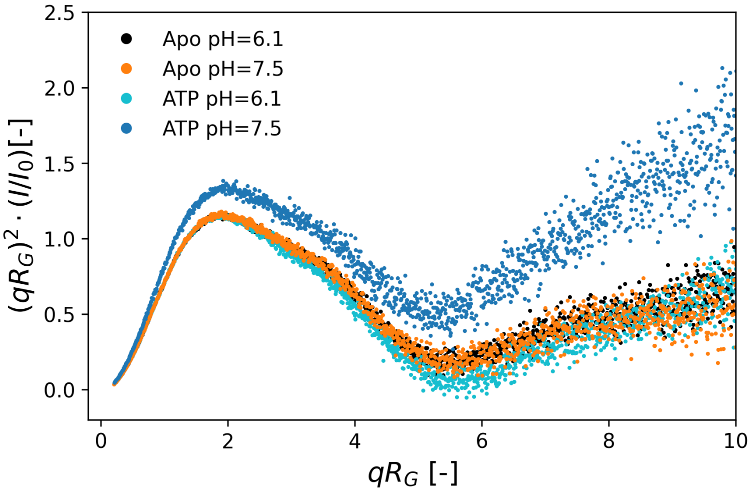


**Figure S1.** Kratky plots obtained from core construct highlight the compact shape of Dhh1 in solution.

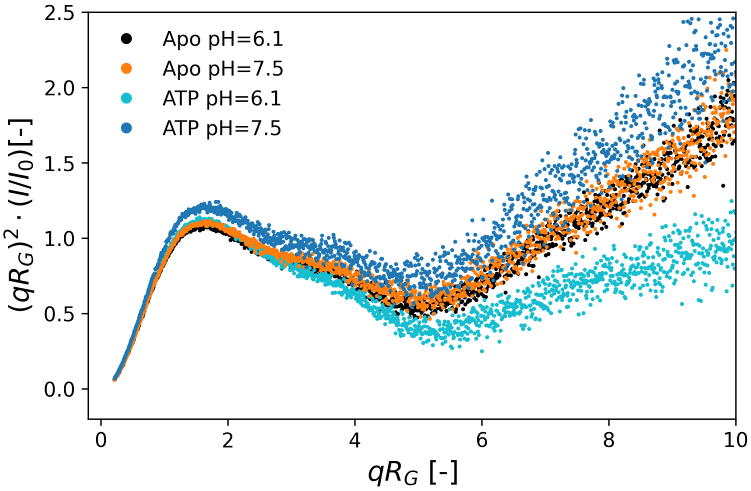


**Figure S2.** Kratky plots obtained from FL construct are compatible with a folded core and the presence of disordered termini.

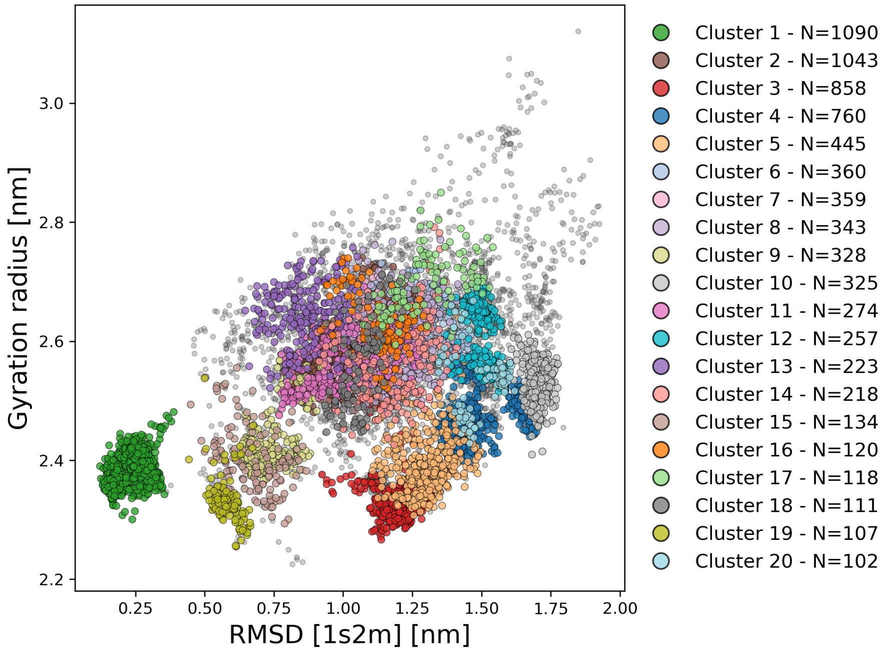


**Figure S3.** Results of conformational clustering plotted on the RMSD/$R_g$ coordinates chosen for the free energy surface. N indicates the population of the cluster. Small light gray points indicate frames that do not own to less populated clusters.

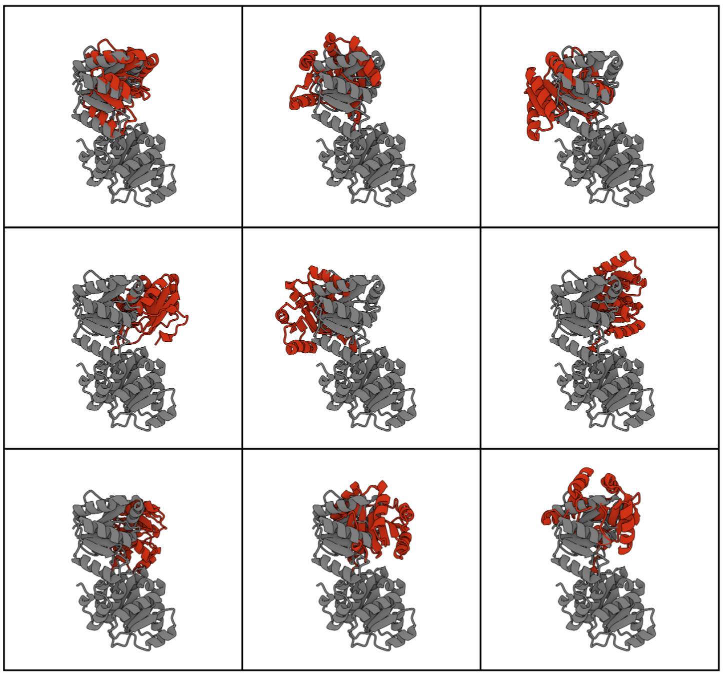

**Figure S4.** The conformations of the centroids (red) of the most populated clusters support that, in solution, the linker is flexible. In gray, the crystallographic structure conformation highlights the different rearrangements of the RecA domains.

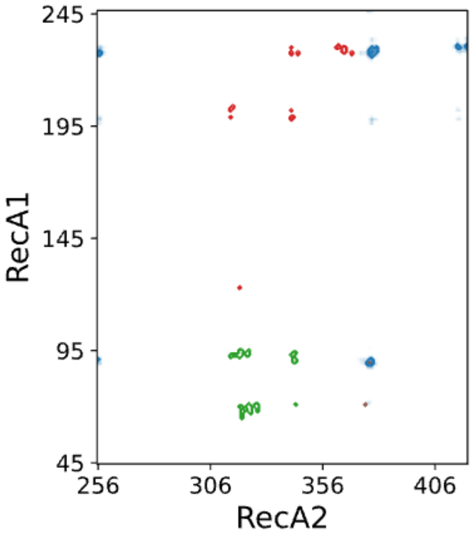


**Figure S5.** The average $C_\alpha$ interdomain contact maps observed in the four most populated clusters (where each colour relates to a given cluster) indicates that the linker flexibility allows different regions of the two RecA domains to interact among them.

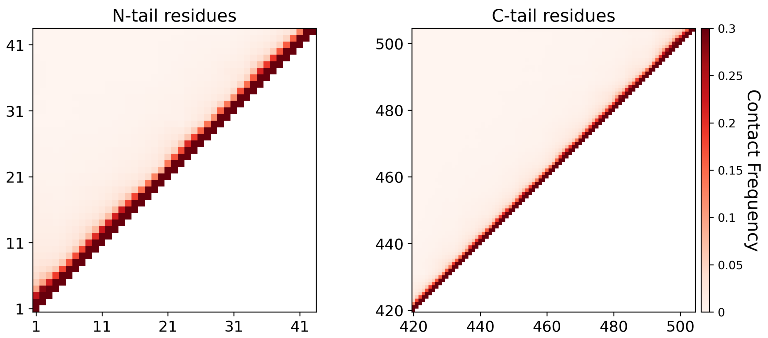


**Figure S6.** Intra-tail contact maps show that none of them display evident interaction patterns. Contact is defined by a cut-off distance of 0.8 nm.

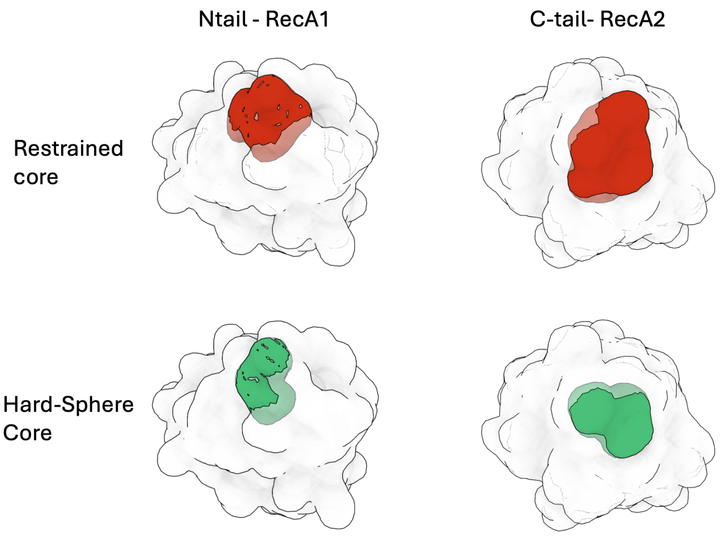


**Figure S7.** Location of the residues that have the highest probability to interact with the N (left) and the C-tail (right) (probability >0.2). There are minimal difference for simulations with restrained core and with hard spheres, with positions topologically close to the attachment point.
